# Comparative transcriptional analysis of *Candida auris* biofilms following farnesol and tyrosol treatment

**DOI:** 10.1101/2023.08.28.555140

**Authors:** Ágnes Jakab, Fruzsina Kovács, Noémi Balla, Csaba Nagy-Köteles, Ágota Ragyák, Fruzsina Nagy, Andrew M Borman, László Majoros, Renátó Kovács

## Abstract

*Candida auris* is frequently associated with biofilm-related invasive infections. The resistant profile of these biofilms necessitates innovative therapeutic options, where quorum sensing may be a potential target. Farnesol and tyrosol are two fungal quorum-sensing molecules with antifungal effects at supraphysiological concentrations. To date there has been no high-throughput comparative molecular analysis regarding the background of farnesol– or tyrosol-related effects against *C. auris* biofilms. Here, we performed genome-wide transcript profiling with *C. auris* biofilms following 75 μM farnesol or 15 mM tyrosol exposure using transcriptome sequencing (RNA-Seq). The analysis highlighted that the number of up-regulated genes (a minimum 1.5-fold increase) was 686 and 138 for tyrosol and farnesol, respectively, while 662 and 199 genes were down-regulated (a minimum 1.5-fold decrease) for tyrosol and farnesol, respectively. The overlap between tyrosol– and farnesol-responsive genes was considerable (101 and 116 overlapping up-regulated and down-regulated genes, respectively). Genes involved in biofilm events, glycolysis, ergosterol biosynthesis, fatty acid oxidation, iron metabolism, and autophagy were primarily affected in treated cells. Farnesol caused an 89.9%, 73.8%, and 32.6% reduction in the calcium, magnesium, and iron content, respectively, whereas tyrosol resulted an 82.6%, 76.6%, and 81.2% decrease in the calcium, magnesium, and iron content compared to the control, respectively. Moreover, the complexation of farnesol, but not tyrosol, with ergosterol is impeded in the presence of exogenous ergosterol, resulting in a minimum inhibitory concentration increase in the quorum-sensing molecules. This study revealed several farnesol– and tyrosol-specific responses, which will contribute to the development of alternative therapies against *C. auris* biofilms.

**Importance:** *Candida auris* is a multidrug-resistant fungal pathogen, which is frequently associated with biofilm related infections. *Candida*-derived quorum-sensing molecules (farnesol and tyrosol) play a pivotal role in the regulation of fungal morphogenesis and biofilm development. Furthermore, they may have remarkable anti-biofilm effects, especially at supraphysiological concentrations. Innovative therapeutic approaches interfering with quorum-sensing may be a promising future strategy against *C. auris* biofilms; however, limited data are currently available concerning farnesol-induced and tyrosol-related molecular effects in *C. auris*. Here, we detected several genes involved in biofilm events, glycolysis, ergosterol biosynthesis, fatty acid oxidation, iron metabolism, and autophagy, which were primarily influenced following farnesol or tyrosol exposure. Moreover, calcium, magnesium, and iron homeostasis were also significantly affected. These results reveal molecular events that provide definitive explanations for the observed anti-biofilm effect; furthermore, they support the development of novel therapeutic approaches against *C. auris* biofilms.

## Introduction

Since its first clinical description, *Candida auris* has grown to represent a serious threat in the healthcare environment, as warranted by the Centers for Disease Control (CDC); in addition, it was assigned to the critical priority group in the fungal priority pathogen list published recently by the World Health Organization (WHO) (1,2). Based on the available literature data, micafungin and amphotericin B have been recommended as the first-line therapy against *C. auris* for adults and infants, respectively (3,4). However, echinocandin resistant-isolate-associated cases have tripled in United States of America in the last two years (5). To further complicate therapy, indwelling medical devices were the source of approximately 90% of *C. auris* candidaemia, indicating that biofilm formation is one of the main predisposing factors of this invasive infection (6,7). In addition, echinocandins – as the first-line therapy – are frequently ineffective for the treatment of these device-related infections. Several data sets are available about the development of resistance to echinocandins following initial administration with these antifungals, particularly in the case of catheter-related infections (8–10).

Quorum sensing is a well-known population density-based communication system through the release and sensing of different quorum-sensing molecules (11,12). In various fungal species, this process plays a pivotal role in the regulation of intra– and inter-species mechanisms such as morphogenesis or virulence (11,12). Farnesol and tyrosol are the two best-described quorum-sensing molecules in the case of *Candida* species. Under physiological conditions, farnesol inhibits the yeast-to-hyphae transition, while tyrosol has the opposite effect in terms of morphogenesis (13,14). The observed potent inhibitory effect of these molecules at supraphysiological concentrations suggests that either farnesol or tyrosol may be a potential part of novel innovative preventive strategies against *Candida* biofilms, including against the *C. auris* sessile community as published previously (15–19). These studies showed that both molecules have a remarkable antifungal effect, interfering with redox homeostasis, virulence, and intracellular microelement contents against planktonic forms of *C. auris*; however, the transcriptome-based biofilm related changes remained to be elucidated (17,18,20).

The present study showed the molecular background of the response to farnesol or tyrosol in *C. auris* biofilm and revealed the transcriptome patterns associated with the observed antifungal effect exerted by these two quorum-sensing molecules. The detailed understanding of quorum sensing molecule-associated molecular mechanisms may open novel innovative therapeutic approaches in the future to overcome this emerging fungal superbug.

## Materials and methods

### Isolate and culture conditions

*C. auris* isolate 12 (NCPF 8973), derived from the South Asian/Indian lineage, was obtained from the National Mycology Reference Laboratory (United Kingdom) (21). The strain was maintained on yeast extract-peptone-dextrose (YPD) solid medium (10 g/l yeast extract [Alfa Aesar, United States of America], 20 g/l mycological peptone [Oxoid, United Kingdom], 20 g/l dextrose, and 20 g/l agar [VWR International Llc, Hungary], pH 5.6). Culturing and biofilm formation were performed in RPMI-1640 (with l-glutamine and without bicarbonate, pH 7.0, and with MOPS; Merck, Budapest, Hungary). Farnesol (Merck Ltd., Budapest, Hungary) was obtained as 3M stock solution, which was diluted to a 30 mM working stock solution in 100% methanol. The working concentration of farnesol were prepared in YPD medium. Drug-free control was supplemented with 1% (vol/vol) methanol. Tyrosol [2-(4-hydroxyphenyl) ethanol] (Merck Ltd., Budapest, Hungary) was prepared as a 0.1 M stock solution in sterile physiological saline.

### Biofilm formation

The *C. auris* isolate was subcultured on YPD agar for 48 hours at 37 °C. Fungal cells were harvested by centrifugation at 3000 × *g* for 5 min and were washed three times with sterile physiological saline. Afterwards, pellets were re-suspended in physiological saline, and the cell density was adjusted to 1 × 10^6^ cells/ml in sterile RPMI-1640 media for each experiment using Burker’s chamber (12,14). The 550 μl suspensions of *C. auris* cells were placed on the bottom of 24-well polystyrene plates (TPP, Trasadingen, Switzerland) to 450 μl RPMI-1640 media and reincubated statically for 24 hours at 37 °C. After the incubation time, the culture medium was aspirated, and non-adherent cells were removed by washing the biofilms with sterile physiological saline. Tyrosol and farnesol in 15 mM and 75 μM concentrations were added to preformed one-day-old biofilms, and then the plates were incubated for 24 hours at 37 °C. Developed biofilms obtained after a further 24 hours of cultivation in the presence and absence of farnesol or tyrosol were scraped from the plate wells with 500 μL of physiological saline and then washed three times with physiological saline (15,17,18). Three biological replicates of biofilm-forming cell suspensions were centrifuged at 3000 g for 10 min at 4 °C, and the pellets were used for RNA extraction and element analysis. Biofilm growth was characterized by dry mass measurement (DCM). The DCM was taken after freeze-drying of the biomass.

### RNA extraction

Similar to our previous studies, total RNA samples were isolated from lyophilized *C. auris* cells (CHRIST Alpha 1-2 LD plus lyophilizer, Osterode, Germany) using Tri Reagent (Merck Ltd. Budapest, Hungary). The quality of RNA was determined using the Eukaryotic Total RNA Nano kit (Agilent, Santa Clara, CA, USA) along with an Agilent Bioanalyzer (18,20).

### Reverse-Transcription Quantitative Real-Time Polymerase Chain Reaction (RT-qPCR) Assays

RT-qPCR was performed to quantify the transcription of 11 selected genes (six up-regulated, *UME6*, *CFL4, BIO2*, *CZF*1, *FAD3*, and *MDR1*; three down-regulated, *PFK1, INO1,* and *POT1;* and two non-differentially expressed genes, *ACT1,* and *ERG9*) selected on the basis of the RNA-Seq experiments. For RT-qPCR, 1 µg of total RNA from each of three independent experiments was digested with DNase I (Merck Ltd. Budapest, Hungary) following the manufacturer’s instructions, and the expression levels of genes were quantified with the Luna® universal one-step RT-qPCR kit (New England Biolabs, Ipswich, MA, USA) with the following cycling parameters: 10 min at 55 °C and 1 min at 95 °C, followed by 40 cycles of 10 s at 95 °C, 10 s at 51 °C, and 20 s at 65 °C. The relative expression of each gene was normalized to that of the *ACT1* (B9J08_000486) gene. Oligonucleotide primers were designed with Oligo Explorer (v.1.1.) and Oligo Analyzer (v.1.0.2) software and are listed in Supplementary Table 1. The relative transcription levels were characterized by the ΔΔCP value. ΔΔCP is the difference between the ΔCPs of the treated and untreated cultures, where ΔCP is the difference between the crossing point of the reference gene and the target gene within a sample (18,20).

### RNA Sequencing

Total RNA was isolated from the farnesol treated, tyrosol treated, and untreated biofilms of *C. auris* isolate 12. Whole RNA sequencing from ∼250 ng of high-quality total RNA (OD_260/280_[≥1.9; RIN value[≥7) was performed at the Genomic Medicine and Bioinformatic Core Facility, Department of Biochemistry and Molecular Biology, Faculty of Medicine, University of Debrecen, Debrecen, Hungary. Libraries were prepared with the NEBNext RNA Sample Preparation kit (New England BioLabs) according to the manufacturer’s protocol. Biofilm samples were sequenced (single-read 75 bp sequencing on an Illumina NextSeq 500 instrument (Illumina, San Diego, California, United States of America) separately. Depending on the sample type, 19–23 million reads per sample (farnesol treated samples), 19–23 million reads per sample (tyrosol treated samples) and 19–23 million reads per sample (untreated samples) were obtained. The FastQC package (www.bioinformatics.babraham.ac.uk/projects/) was used for quality control. Reads were aligned to the genome of *C. auris B8441*, retrieved from the Candida Genome Database (CGD) (www.candidagenome.org) with the HISAT2 algorithm combined with SAMtools (22). The successfully aligned reads varied between ≥92% (farnesol treated samples), ≥92% (tyrosol treated samples), and ≥92% (untreated samples). Downstream analysis was performed using StrandNGS software (www.strand-ngs.com). BAM files were imported into the software, and the DESeq algorithm was used for normalization. A moderated *t*-test was used to determine differentially expressed genes between conditions.

### Evaluation of transcriptome data

The Candida Genome Database platform (www.candidagenome.org) with default settings was used to characterize the up– and down-regulated gene sets. Only hits with a corrected *p* value < 0.05 were regarded as significantly enriched (Table S2 and S3). Enrichment of selected gene groups in the up– and down-regulated gene sets was also studied with the Fisher’s exact test function of the R project (www.R-project.org/) (Table S3).

#### The following gene categories were examined

The “Virulence-associated genes” are known as putative genes involved in the genetic regulation of *C. albicans* virulence properties according to previously published classifications (23–25).

The “Metabolic pathway-associated genes” include all genes related to ergosterol, carbohydrate, and fatty acid biochemical pathways based on the pathway databases (http://pathway.candidagenome.org/).

The “Metal metabolism-associated genes” group involved in manganese, calcium, magnesium, iron, zinc, and copper homeostasis genes by *C. albicans* were collected by the method of Fourie *et al*. (2018) and Gerwien *et al*. (2018) (26,27).

“Autophagy-related genes” were collected from the Candida Genome Database (www.candidagenome.org).

### Intracellular metal content measured by inductively coupled plasma optical emission spectrometry (ICP-OES) in *Candida auris* biofilms

The selected intracellular element (Fe, Ca, and Mg) contents of the lyophilized biomass were determined by inductively coupled plasma optical emission spectrometry (ICP-OES; 5110 Agilent Technologies, Santa Clara, CA, USA) following atmospheric wet digestion in 3 ml of 65% HNO_3_ and 1 ml of 30% H_2_O_2_ in glass beakers. The metal contents of the samples were normalized by DCM as described by Jakab *et al.* (2021) (20). The metal contents of the dry biomass were determined in triplicate, and mean ± standard deviation values were presented.

### Ergosterol-binding assay

To determine the binding of farnesol or tyrosol to the ergosterol present in *C. auris* cell membranes, an ergosterol binding assay was performed as described by Ramesh *et al.* (2023) (28). Briefly, ergosterol (Merck, Budapest, Hungary) was dissolved in dimethyl-sulfoxide (DMSO). The prepared ergosterol solution was then pipetted to RPMI-1640 in 100 and 200 mg/l final concentrations. The minimum inhibitory concentration (MIC) values of farnesol or tyrosol against *C. auris* were determined in RPMI-1640 according to the recommendations proposed by the Clinical Laboratory Standards Institute M27-A3 protocol with and without media supplemented with ergosterol (29). The concentrations tested ranged from 0.585 to 300 μM for farnesol and from 0.058 to 30 mM for tyrosol, with 100 and 200 mg/l of ergosterol in RPMI-1640. MICs were determined as the lowest concentration that caused at least 50% growth decrease compared with the untreated control cells. The changes in MIC values with and without of added ergosterol were determined to conclude the ergosterol-binding ability of farnesol and tyrosol.

## Data availability

Regarding the *C. auris* isolate tested, the Whole Genome Shotgun project has been deposited in DDBJ/ENA/GenBank under the accession JANPVY000000000. Transcriptome data have been deposited in NCBI’s Gene Expression Omnibus (GEO; http://www.ncbi.nlm.nih.gov/geo/) and are accessible through GEO Series accession number GSE233427 (https://www.ncbi.nlm.nih.gov/geo/query/acc.cgi?acc=GSE233427).

## Results

### Genome-wide transcriptional changes for *C. auris* biofilms

Long-term transcriptional responses were studied for *C. auris* biofilms in three different experimental settings: (i) untreated control *C. auris* biofilm, (ii) farnesol-exposed *C. auris* biofilm, and (iii) tyrosol-treated *C. auris* biofilm. In these experiments, one-day-old biofilms were supplemented with 75 μM farnesol or 15 mM tyrosol and samples were collected after 24 hour-long exposures. Reproducible relationships between RNA-Seq results were confirmed by principal component analysis (Figure S1). The effects of quorum-sensing molecules on the transcriptomes are summarized in Figures 1A-D and 2.

**Figure 1.**
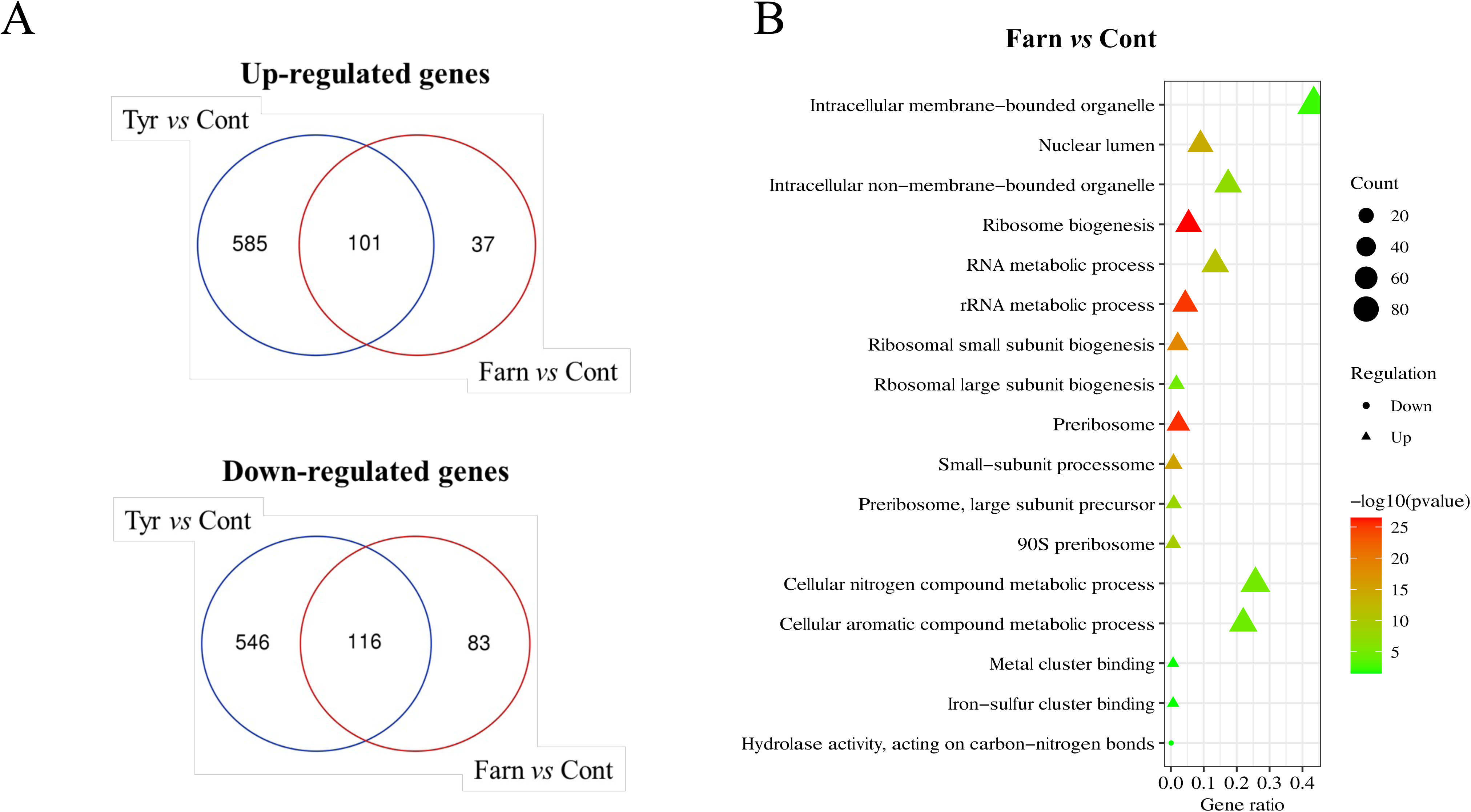
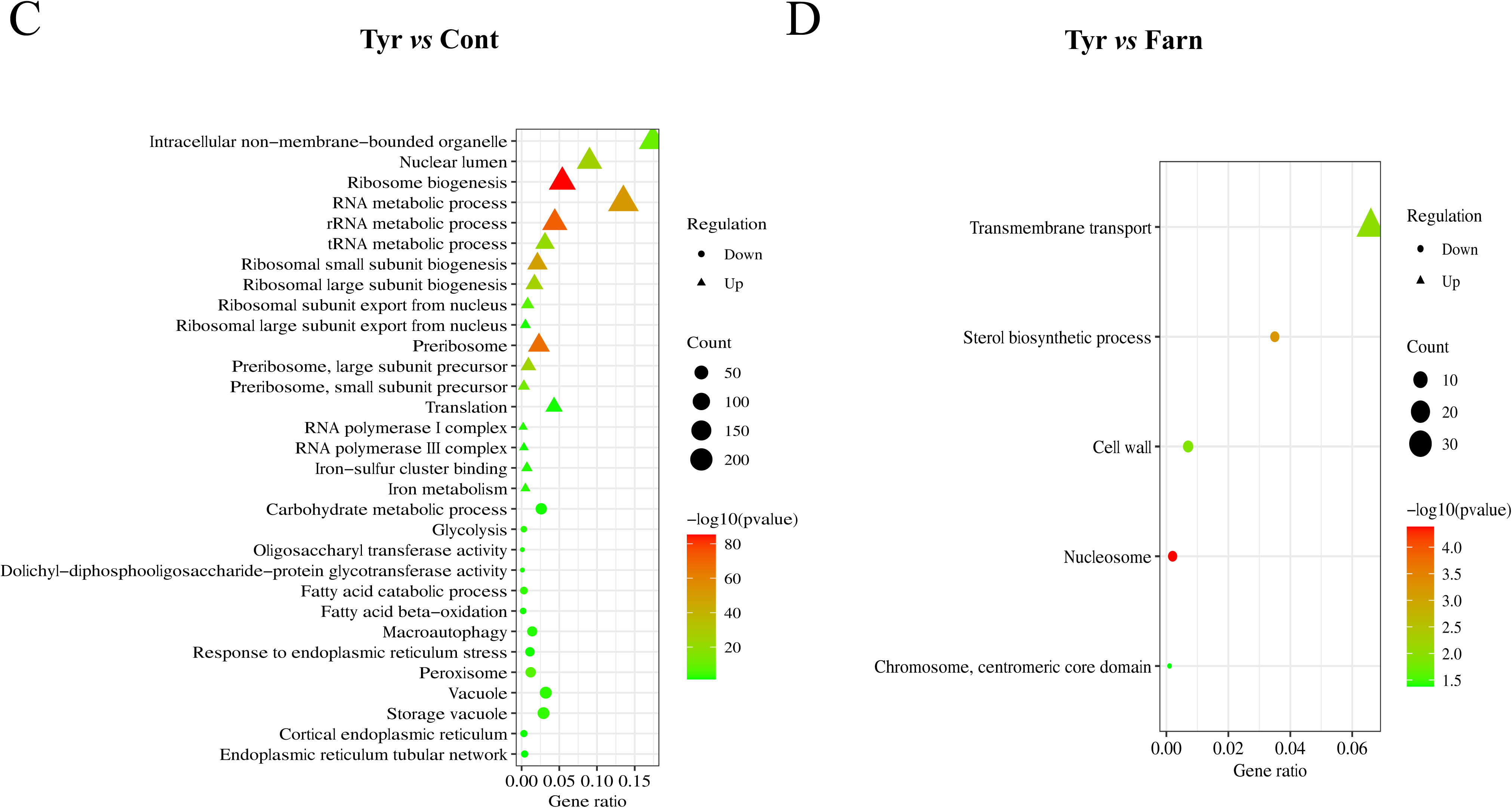
S**u**mmary **of RNA-Seq data and gene enrichment analyses.** (A) The effects of tyrosol (Tyr *vs*. Cont) and farnesol (Farn *vs.* Cont) treatment to the transcriptomes are depicted in the Venn diagrams. (B-D) Bubble charts of Gene Ontology (GO) terms of Candida Genome Database (http://www.candidagenome.org/cgi-bin/GO/goTermFinder) generated by different expression genes. Bubble charts represent up(Δ)– and down(•)-regulated genes belonging to gene groups farnesol treated versus untreated (B), tyrosol treated versus untreated (C) and farnesol treated versus tyrosol treated (D) comparisons where the enrichment was significant (p < 0.05). The color of bubble means the significance of the corresponding GO pathway (in green color, low *p* values; in red color, high *p* values). As well, the size of bubble means the number of different expression genes in this pathway. Only the differentially expressed genes (corrected *p* value of <0.05) exhibiting more than 1.5-fold increase or decrease in their transcription are shown. The full list of the gene groups is available in Supplementary Tables S2 and S3.

**Figure 2.**
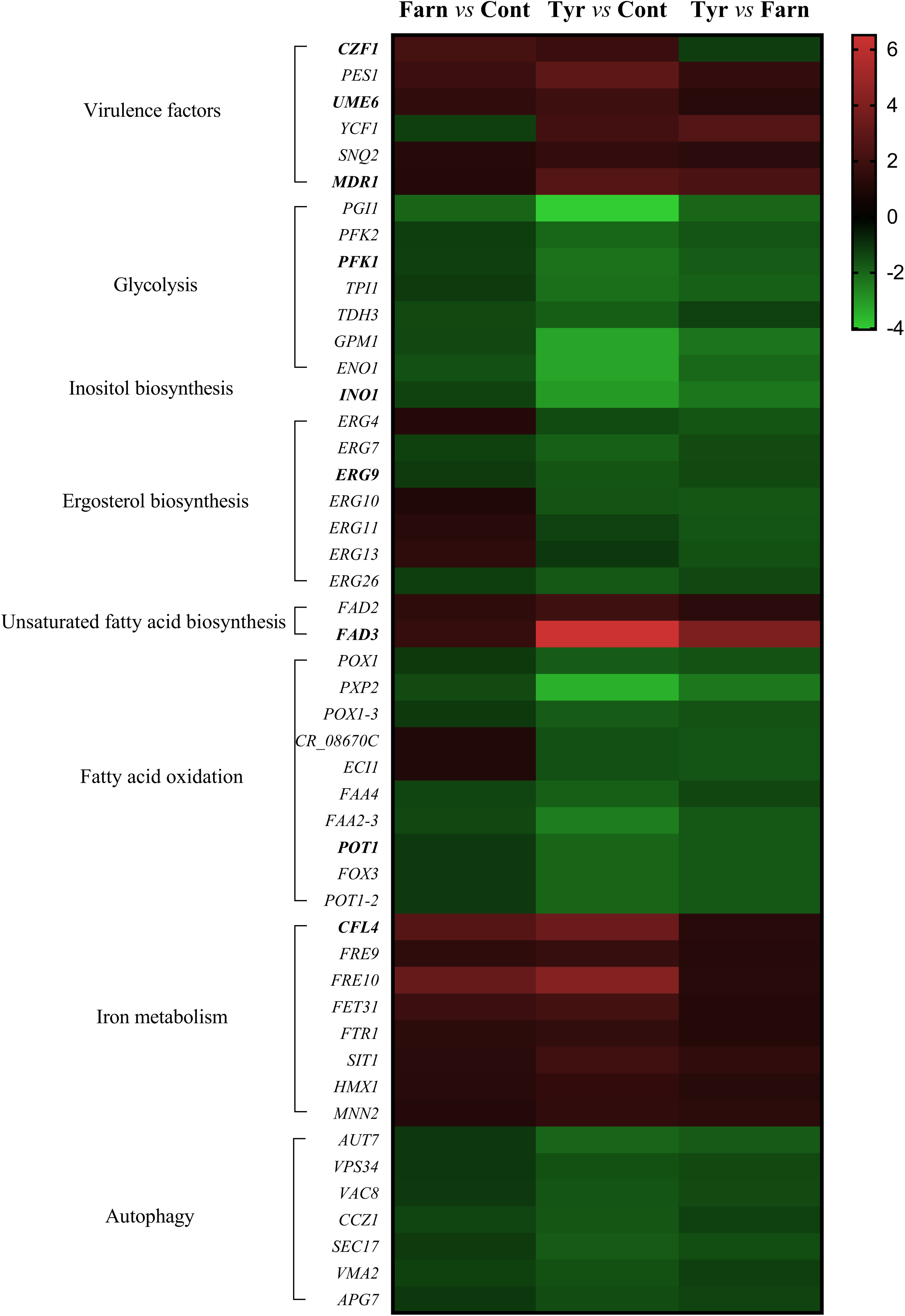
T**h**e **effects of quorum sensing compounds, farnesol and tyrosol, on the expression of selected genes of *C. auris* biofilms.** The heat map demonstrates the expression profiles of representative genes according to the color scale that indicates gene expression changes in FC units. Supplementary Table S3 summarizes the data that were used for the construction of the heat map.

Tyrosol-related effects were more pronounced on *C. auris* biofilms compared to untreated control sessile cells. The number of up-regulated genes were 686 and 138 for tyrosol and farnesol, respectively; while 662 and 199 genes were down-regulated for tyrosol and farnesol, respectively (Figure 1A). The overlaps between tyrosol– and farnesol-responsive genes were considerable (101 and 116 overlapping up-regulated and down-regulated genes, respectively); however, the transcription of several genes changed exclusively in response to tyrosol exposure (the number of up-regulated and down-regulated tyrosol-responsive genes was 585 and 546, respectively (Figure 1A).

The fold change obtained using RNA-Seq was compared with relative transcription levels (ΔΔCP) derived from RT-qPCR analysis. Selected genes of interest are shown in Supplementary Table S1. The similarity between the transcription levels obtained from the two methods indicates high consistency between the analytical data. Supplementary Table S4 indicates a good correlation between RNA-Seq and RT-qPCR data with a correlation coefficient (r) of 0.89 (Farnesol *vs.* Control) and 0.95 (Tyrosol *vs.* Control). The possible physiological background of the transcriptional changes for up– and down-regulated genes was further characterized using gene set enrichment analyses (Figure 1B-D, Tables S2 and S3), and selected changes are illustrated in a heat map (Figure 2).

### Farnesol exposure shows a moderate transcriptomic effect

Based on our transcriptomic data, genes involved in biofilm formation (*CZF1*, *UME6*, and *TYE7* transcription factors; and *PES1*, encoding a key enzyme specific to regulation of the hyphae-to-yeast transition), iron-sulfur cluster binding (*RLI1*, *ISA1*, *BIO2*, *SDH2*, *DRE2*, and *LEU1*), iron uptake (*FET31*, multicopper oxidase; *CCC1*, ferrous iron transporter; *CFL4* and *FRE10,* ferric reductases), as well as in ribosome biogenesis (50 genes), ribosomal small subunit biogenesis (29 genes), ribosomal large subunit biogenesis (14 genes), RNA metabolic process (54 genes) and rRNA metabolic process (45 genes) were enriched in the up-regulated gene set (Figures 1 and 2, Tables S2 and S3). Up-regulation of *UME6*, *CZF1*, *BIO2,* and *CFL4* under farnesol exposure was confirmed by RT-qPCR data (Tables S4).

### Tyrosol treatment led to a considerable reprogramming of gene transcription in *C. auris* **biofilm**

Transcripts of biofilm-formation genes (*CZF1*, *UME6*, and *TYE7* transcription factors; and *PES1*) were also activated by tyrosol treatment (Figure 2, Table S3). Furthermore, significant up-regulation was observed in the case of the following genes: putative ABC transporters (*MDR1*, *YCF1,* and *SNQ2*), unsaturated fatty acid biosynthetic process (*FAD2* and *FAD*3 encoding for delta-12 and omega-3 fatty acid desaturases), iron homeostasis (*CFL4*, *FRE9*, and *FRE10*, ferric reductases; *FET31*, multicopper oxidase; *FTR1*, iron permease; *SIT1*, ferrichrome siderophores transporter; *HXM1*, heme oxygenase; *MNN2* and *CCC1*, iron transporters), iron-sulfur cluster binding (15 genes, e.g., *RLI1*, *ECM17*, *YAH1*, *ISA1*, *LYS4*, *BIO2*, *ELP3*, *SDH2*, *DRE2*, and *LEU1*), as well as ribosome biogenesis (175 genes), ribosomal small subunit biogenesis (84 genes), ribosomal large subunit biogenesis (57 genes), RNA metabolic process (241 genes), rRNA metabolic process (147 genes), tRNA metabolic process (74 genes), RNA polymerase I complex (8 genes), RNA polymerase III complex (9 genes), and translation (52 genes) (Figures 1 and 2, Tables S2 and S3).

On the other hand, ergosterol biosynthetic process (*ERG4*, *ERG7*, *ERG9*, *ERG10*, and *ERG26*), phospholipid binding (26 genes), carbohydrate metabolic process (36 genes), inositol metabolic process (*INO1*, and CR_08330W), trehalose metabolism (*TPS1*), carbohydrate catabolic process (17 genes), glycolysis (*PGI1*, *PFK2*, *PFK1*, *TPI1*, *TDH3*, *GPM1*, and *ENO1*), maltose degradation *(MAL2*, C5_04940W, *GDB1*, and C5_04940W), fatty acid metabolic process (20 genes), fatty acid catabolic process (11 genes), fatty acid beta-oxidation (*POX1*, *PXP2*, *POX1-3*, CR_08670C, *ECI1*, *FAA4*, and *FAA2-3*), glyoxylate cycle (*MLS1*, and *MDH1-3)*, glutamate catabolic process (*GAD1*, *UGA11*, and *UGA2*), copper uptake (*CTR1,* copper transporter; *CCS1,* copper chaperone) zinc metabolism (*PRA1*, surface protein; *CSR1*, transcription factor), as well as macroautophagy (25 genes), and response to endoplasmic reticulum stress (20 genes) were enriched in the down-regulated gene set (Figures 1 and 2, Tables S2 and S3).

Moreover, tyrosol exposure significantly decreased the transcription of 28 peroxisomal genes, 42 vacuolar genes, 37 genes of the cell cortex, including 9 genes of the cortical endoplasmic reticulum and 20 genes of the cortical actin cytoskeleton, as well as 9 genes of the endoplasmic reticulum tubular network in the cellular component-related term (Figures 1 and 2, Tables S2 and S3). It is noteworthy that tyrosol treatment caused a significant increase in transcription of *UME6*, *CZF1*, *FAD3*, *BIO2*, *CFL4*, and *MDR1* based on the RT-qPCR measurements. In addition, down-regulation of *PFK1*, *INO1*, and *POT1* was also supported by RT-qPCR (Tables S4).

The obtained data indicated that tyrosol exposure significantly increased the transcription of 30 transmembrane transport-related genes and decreased the expression of 4 ergosterol biosynthetic process (*ERG4*, *ERG10*, *ERG11*, and *ERG13*)-related genes compared to farnesol treatment (Figures 1 and 2, Tables S2 and S3).

### Quorum-sensing molecules significantly influenced the metal content of one-day-old *C. auris* biofilm

Quorum-sensing molecules decreased the dry cell mass and metal contents significantly compared to untreated control biofilms (*p* < 0.01), as presented in Table 1. A significant decrease was detected in the DCM of farnesol– and tyrosol-treated biofilms (0.53 ± 0.165 g/l and 0.4 ± 0.16 g/l for farnesol and tyrosol, respectively) compared to untreated cells (1.37 ± 0.35 g/l). Furthermore, farnesol and tyrosol exposures significantly influence the calcium (319.37 ± 234.80 mg/kg and 551.75 ± 441.83 mg/kg), magnesium (695.78 ± 111.91 mg/kg and 618.65 ± 40.75 mg/kg), and iron (240.34 ± 118.39 mg/kg and 67.17 ± 15.84 mg/kg) contents of *C. auris* biofilms as compared to controls (3170.7 ± 82.8 mg/kg for calcium, 2648.36 ± 35.05 mg/kg for magnesium and 356.32 ± 45.62 mg/kg for iron, respectively) (Table 1).

**Table 1.** Effects of quorum sensing molecules significantly influences the metal contents of *Candida auris* biofilms.

| Culture | Dry cell mass (DCM) (g/l) | Metal contents/treatment (mg/kg) (mean ± SD<sup>a</sup>) | | |
| --- | --- | --- | --- | --- |
|  |  | Ca | Mg | Fe |
| **Control cultures** | 1.37±0.35 | 3170.7±82.8 | 2648.36±35.05 | 356.32±45.62 |
| **+ 75 μM Farnesol** | 0.53±0.165** | 319.37±234.80** | 695.78±111.91** | 240.34±118.39 |
| **+ 15 mM Tyrosol** | 0.4±0.16** | 551.75±441.83** | 618.65±40.75** | 67.17±15.84** |

<sup>a</sup> Mean values ± standard deviations (SD) calculated from three independent experiments are presented.
The asterisks indicate significant differences calculated by Student t test comparing untreated control and farnesol or tyrosol-treated cultures as follows: \*\* $p < 0.01$ .

### Ergosterol-binding assay

The ability of farnesol or tyrosol to cause membrane destabilization can be identified by its ability to interfere with exogenous ergosterol added to the *C. auris* suspension in standard microdilution assay. In the presence of exogenous ergosterol at 100 and 200[mg/l, the MIC of farnesol increased 4-fold, from 75 to 300 μM for *C. auris*. In the combination of tyrosol and ergosterol, the MIC values were 30 mM in the presence or absence of ergosterol. These results indicate that farnesol but not tyrosol may exert its activity in whole or in part by binding to membrane ergosterol.

## Discussion

In the past decade, *C. auris* has caused multiple outbreaks worldwide, which have frequently been associated with extensive biofilm production on indwelling devices, further complicating the already challenging treatment (5–7,30). Previous studies revealed that innovative anti-biofilm strategies interfering with quorum sensing may effectively attack this hard-to-treat sessile pathogen (31,32). Fungal quorum-sensing molecules (farnesol or tyrosol), especially at supraphysiological concentrations, have remarkable antifungal and drug potentiator effects against several *Candida* species (15–20). In the case of planktonic *C. auris* cells, the molecular and physiological background of farnesol-related effects were described; however, the biofilm-specific molecular and physiological events following farnesol or tyrosol exposure remained to be elucidated so far (20). It is noteworthy that previously performed differential expression analysis demonstrated that the *C. auris* planktonic and biofilm transcriptome differ significantly (33). Therefore, the planktonic findings could not be directly extrapolated to biofilms. In this study, we performed comprehensive comparative transcriptomic profiling to significantly expand the list of genes affected by farnesol or tyrosol and those physiological processes that will be able to support the development of effective therapies against *C. auris* biofilms.

Our comparative transcriptomic data show a significant up-regulation in *CZF1* and *UME6* genes following both farnesol and tyrosol exposure. Similarly up-regulated was *TYE7*, which is the major transcriptional regulator of glycolysis genes in *C. albicans* that attaches the promoters of genes related to glycolysis such as *PFK1* and *PFK2* encoding subunits of phosphofructokinase (34). This enzyme irreversibly converts fructose-6-phosphate into fructose-1,6-bisphosphate, which is a pivotal regulatory step in glycolysis (34,35). Furthermore, it acts as a negative regulator of hypoxic filamentation (36). Despite the overexpression of *TYE7*, several key genes in glycolysis were significantly down-regulated (*PGI1*, *PFK1*, *PFK2*, *TPI1*, *TDH3*, *GPM1*, *ENO1*), especially under tyrosol exposure. The opposite pattern was reported in *C. parapsilosis* planktonic cells, where exogenous tyrosol treatment shifted metabolism toward glycolysis (18). Overexpression of *CZF1* protein stimulates filamentation; moreover, *CZF1* gene deletion is associated with negative effects on hyphae filamentation. Similar *CZF1* up-regulation was observed in case of *C. parapsilosis* planktonic cells following tyrosol exposure; however, Jakab et al (2019) did not observe higher rates of adherence and biofilm-forming ability in the presence of this quorum-sensing molecule (18). Gene of *UME6* is also important in terms of hyphal extension. In addition, Ume6 protein has a pivotal role in the expression of *HWP1*, *ECE1*, *ALS3*, and *HCG1*, which are associated with the filamentation (35,36). We hypothesize that the observed up-regulation of *CZF1* and *UME6* is a compensatory response of fungi to maintain the biofilm structure, because both farnesol and tyrosol exposure significantly decreased the level of two bivalent cations – magnesium and calcium – which play a critical role in biofilm development.

Previous studies suggest that magnesium triggers the growth of filamentous forms in *C. albicans* and in *Trichosporon asahii* (37,38). Furthermore, magnesium uptake has an effect on mitochondrial distribution, the production of lipid droplets, and vacuolar growth, which contribute to promotion of hyphal growth and directly to biofilm formation (38). Hans *et al.* (2019) showed that magnesium deprivation impedes the metabolic flexibility of *C. albicans* (39). In our study, several glycolysis, gluconeogenesis, and fatty acid oxidation-related genes were down-regulated, especially for tyrosol treatment, which were associated with the reduced growth rate and the significantly decreased dry cell mass of sessile cells. A previous study revealed that magnesium chelation and its lower level leads to the potentiation of membrane-targeting antifungal drugs, which was confirmed previously between farnesol and triazoles against *C. auris* biofilms (17,39). In addition, the decreased magnesium content inhibited potential virulence traits, including biofilm formation, morphological transition, and adherence to epithelial cells; moreover, it significantly influences membrane homeostasis with remarkable changes in ergosterol synthesis-related genes, as confirmed in this study (39). Aside of the magnesium content, both farnesol– and tyrosol-treated biofilms showed a decreased calcium level. Previous results demonstrated that calcium supplementation could increase the length of fungal cells grown for *T. ashaii*, *Cryptococcus neoformans*, and *C. albicans* because calcium regulates both actin polymerization and microtubule polymerization; thus, it has a remarkable direct effect on biofilm development (40,41). In accordance with these studies, beside of decreased calcium level, tyrosol treatment significantly down-regulated the transcription of several genes, which influence the actin filament organization, actin cortical patch, cortical cytoskeleton and cortical actin cytoskeleton. Presumably, the simultaneous reduction of these two crucial bivalent cations (magnesium and calcium) may explain the previously documented anti-biofilm effect exerted by farnesol or tyrosol.

Tyrosol treatment significantly decreased the iron content of biofilms, which were associated with several up-regulated iron homeostasis-related gene groups (e.g., ferric reductases, multicopper oxidases, iron permeases). Although farnesol exposure resulted in a similar pattern in the transcription level of these genes, the observed changes did not coincide with a significantly decreased iron content. Nevertheless, previously published planktonic *C. auris* transcriptomic data showed that farnesol treatment down-regulated the transcription of iron homeostasis-related genes, which were associated with a significant reduction of the iron concentration (20). It is noteworthy that iron deprivation does not influence the biofilm-forming ability of *C. albicans* (42). Nonetheless, the decreased iron content enhances the membrane fluidity of *Candida* cells, influencing its susceptibility to membrane-active antifungal agents (43).

Considering the results derived from transcriptome analysis, intracellular metal content determination and ergosterol-binding assay, the examined fungal quorum-sensing molecules appears to impact the fungal cell membrane structure. Our ergosterol binding assay shows that farnesol is highly bound to the ergosterol, which presumably changes the conformational properties of ergosterol, influencing the membrane characteristics; nevertheless, further structure-based confirmatory experiments are needed to justify this hypothesis. Tyrosol could also influence certain membrane characteristics. Tyrosol treatment significantly enhanced the transcription of *FAD2* and *FAD3* genes encoding for fatty acid desaturases involved in poly unsaturated fatty acid synthesis. Riekhof *et al*. (2014) demonstrated a similar pattern in *FAD2*/*FAD3* transcription following phosphate starvation in fungi (44). The overexpression of these desaturases may increase the tolerance of fungal cells to environmental stress.

Another remarkable tyrosol-induced membrane related effect was the down-regulation of several ergosterol synthesis-associated genes including *ERG4*, *ERG7*, *ERG9*, *ERG10*, and *ERG26*. The down-regulation of these genes may alter the membrane’s permeability and may influence its fluidity. Regarding farnesol, Dizova *et al.* (2017) showed that farnesol exposure (200 μM) down-regulated the *ERG9*, *ERG11*, and *ERG20* genes in *C. albicans* (45). Furthermore, Jakab *et al*. (2021) reported that the presence of 75 μM farnesol decrease the transcription of *ERG6* gene in *C. auris*, which might enhance the passive diffusion of farnesol; moreover, the decreased *ERG6* content produces higher susceptibility to oxidative stress and impairs thermotolerance (20). Surprisingly, farnesol did not cause any relevant change in the transcription of central ergosterol biosynthesis-related genes in this study. Aside from *ERG* genes, *INO1*, encoding inositol-1-phosphate synthase, was also down-regulated following tyrosol exposure. Interestingly, in the case of planktonic *C. auris* cells, farnesol reduces the transcription of this gene (20).

With respect to autophagy-related genes, tyrosol exposure caused a significant decrease in the transcription of *C1_00430W*, *AUT7*, *VPS34*, *C4_01790W*, *VAC8*, *CCZ1*, *C7_03860W*, *SEC17*, *VMA2*, and *APG7*, whereas the transcription level of *SPO72* was increased. Macroautophagy is an evolutionarily conserved dynamic pathway that functions primarily in a degradative manner. Macroautophagy has a pivotal role in maintenance of cellular homeostasis; however, either under-activated or over-activated macroautophagy can remarkably compromise cell physiology, leading to cell death (46).

This is the very first study analyzing the global changes in gene transcription of *C. auris* biofilms in a comparative manner following farnesol and tyrosol exposure. Nevertheless, one major limitation should be highlighted. The *C. auris* isolates are classified into five different clades (47). These lineages differ by several thousand single nucleotide polymorphisms. In this study, we examined only one isolate from one clade (South Asian lineage). Nevertheless, among all clades, the South Asian clade contains the highest percentage of multidrug-resistant isolates (48). Thus, the obtained results are relevant in terms of overcoming biofilms formed by multidrug-resistant isolates from the South Asian clade. Taken together, our data give a novel insight into the genome-wide transcriptome changes caused by farnesol and tyrosol exposure in the metal content of biofilms, metabolic regulation, and membrane-related alterations. However, further mutant-based *in vitro* and *in vivo* investigations are needed to fully understand the complete mechanism of these two quorum-sensing molecules in the *C. auris* sessile community.

## Acknowledgements

R. Kovács was supported by the Janos Bolyai Research Scholarship of the Hungarian Academy of Sciences (BO/00127/21/8). This research was supported by the Hungarian National Research, Development and Innovation Office (NKFIH FK138462). R. Kovács was supported by the UNKP-22-5-DE-417 New National Excellence Program of the Ministry for Innovation and Technology from the Source of the National Research, Development and Innovation Fund.

## Legends to the Figures

**Supplementary Figure 1**: Principal component analysis of the transcriptome data (A) and Clusters (B). Symbols represent untreated control (Cont) 75 μM farnesol (Farn) and 15 mM tyrosol exposure (Tyr) cultures. Analyses were performed with the StrandNGS software using default settings.

**Supplementary Table 1**: Oligonucleotide primers used for RT-qPCR analysis.

**Supplementary Table 2**: Results of the gene set enrichment analysis.

Significant shared GO terms (p < 0.05) were determined with Candida Genome Database Gene Ontology Term Finder (http://www.candidagenome.org/cgi-bin/GO/goTermFinder). Biological processes, molecular function and cellular component categories are provided.

**Supplementary Table 3**: Transcription data of selected gene groups.

Part 1: Genes involved in genetic control of *Candida auris* virulence.

Part 2: Genes involved in metabolism.

Part 3: Genes involved in ergosterol and fatty acid metabolic process.

Part 4: Genes involved in metal metabolism.

Part 5: Genes involved in autophagy.

The systematic names, gene names, gene orthologs in *Candida albicans* and the features (putative molecular function or biological process) of the genes are given according to the Candida Genome Database (http://www.candidagenome.org). Up– and downregulated gene were defined as differentially expressed genes (corrected *p* value < 0.05) where log_2_(FC)>0.585 or log_2_(FC)< –0.585, respectively, and FC stands for fold change ratios (tyrosol treated *vs*. untreated) and are marked with red and blue colour. Results of gene enrichment analysis (Fisher’s exact test) are also enclosed.

**Supplementary Table 4**. Overview of RT-qPCR assays

RNA-Seq data are presented as FC values, whereby FC is abbreviation of “fold-change”. Relative transcription levels (ΔΔCP) were quantified with ΔΔCP = ΔCP_control_ –ΔCP_treated_. CP values stand for the qRT-PCR cycle numbers of crossing points. The *ACT1* (B9J08_000486) was used as reference gene. RT-qPCR data are presented as mean ± SD calculated from three independent measurements. Significantly higher or lower than zero ΔΔCP values (up-regulated or downregulated gene) are marked with red and blue colors, respectively (Student’s t-test, *p* < 0.05, n = 3). Diagrams demonstrate the correlation between RT-qPCR and RNA-Seq data.

