## Supplementary figures and images for "Comparative transcriptional analysis of *Candida auris* biofilms following farnesol and tyrosol treatment"

### Supplemental Figure 1

Figure S1 Principal component analysis of RNAseq data

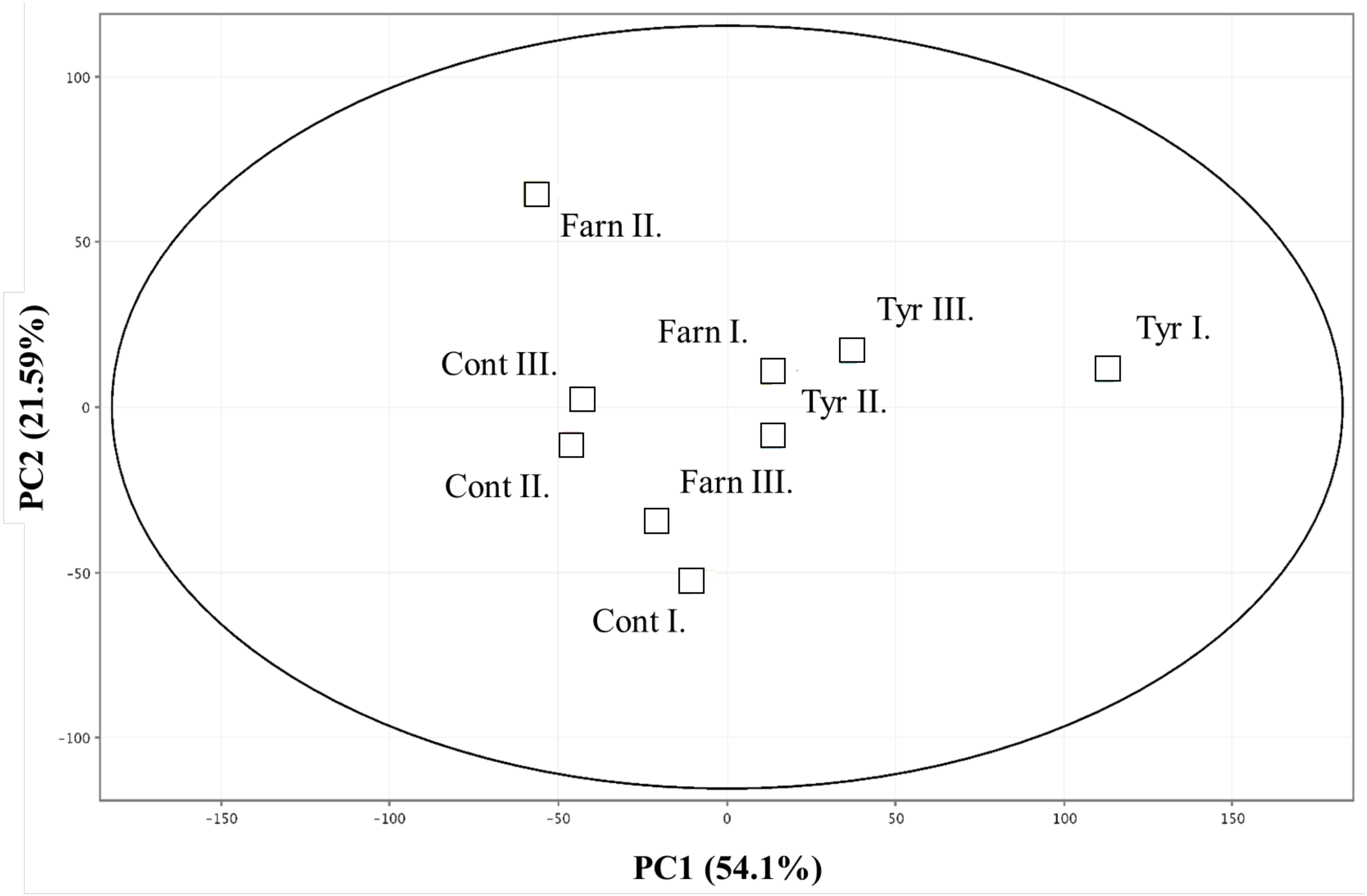
